# Drought resilience of long term dry-farmed grapevines *(Vitis vinifera L.)*

**DOI:** 10.1101/2021.04.07.438886

**Authors:** Vinay Pagay, Tarita S. Furlan, Catherine M. Kidman, Dilrukshi Nagahatenna

**Author notes:** *****Corresponding author. *Email address:* (V. Pagay).

## Abstract

We explored the long-term drought resilience of field-grown unirrigated (or dry-grown) grapevines of pre-clonal origin from shallow (SR) and deep (DR) soils representing low and high soil water availability, respectively, in a warm, Mediterranean climate. Despite lower soil moisture available to SR vines, both deep and shallow-rooted vines had similar vine water status, based on measurements of midday stem water potential (Ψ_*s*_), and leaf net photosynthesis (*A*_*n*_). Due to the lower stomatal conductance (*g*_*s*_), SR had higher intrinsic water use efficiency (*WUE*_*i*_) than DR, however the carbon isotope ratio (*δ*^13^*C*) of the fruit at harvest was similar between the two groups. Our observations suggest a degree of drought adaptation in the SR vines resulting from multi-decadal cyclical droughts. Overall, we demonstrate that pre-clonal Cabernet Sauvignon grapevines dry-grown in shallow soils have an enhanced resilience to drought compared to dry-grown vines in deep soils. This study has implications for selection of crop genetic material in a changing climate.

## 1. Introduction

Climate change has increased the pressure on freshwater resources globally, particularly in arid and semi-arid regions, the latter where a significant proportion of grapevines are commercially grown. In Australia, for example, the availability of fresh water is predicted to decline; climate change data indicated a 10-20% decrease in rainfall in Australia between 1997 and 2013 (relative to the reference period, 1900-2013), primarily in the cooler months between May and July when most of the annual rainfall is received, with large impacts on groundwater levels (CSIRO and BOM, 2015). The predictions for South Australia to 2050 indicate a similar scenario with yearly rainfall to decrease between 5% and 15%, and up to 21% during spring (DEW, 2018). Coupled with extreme events during the growing season, e.g. heatwaves, the water loss from vineyards through both soil evaporative losses and vine transpiration rates will very likely increase.

Soil moisture deficits and consequent vine water stress result in several physiological responses spanning both vegetative and reproductive processes. These responses have been extensively documented, and have formed the basis of several seminal and review papers (Chaves et al., 2010; Lovisolo et al., 2016; Schultz and Stoll, 2010). In red grape production where wine quality is important, excess irrigation may negatively impact on grape primary (e.g. sugars) and secondary (e.g. flavonoids) metabolites that contribute to wine quality (McCarthy, 1997; Salon et al., 2005). Therefore, optimizing irrigation scheduling to maintain a level of water stress in the vine that enhances grape berry/wine quality while simultaneously maximizing vineyard water use efficiency (WUE = yield/total water incident on vineyard) is of paramount importance to vineyard managers seeking to sustainably produce high-quality fruit.

Several vineyard management strategies to achieve higher WUE have been proposed including deficit irrigation (Chaves et al., 2010; Dry et al., 2001), use of specific scions (Chaves et al., 2010) and rootstocks (Zhang et al., 2016), use of mulches and cover crops (Beslic et al., 2015), reduced use of nitrogen-based fertilizers and specific vine training systems (van Leeuwen and Destrac-Irvine, 2017) and, more recently, the use of soil and crop water sensing technologies to schedule irrigation more precisely (Pagay and Kidman, 2019; Rienth and Scholasch, 2019; Yu and Kurtural, 2020). Another approach of increasing WUE that has received less attention is dry-faming or dry-growing grapevines, which refers to growing grapevines without supplemental irrigation, i.e. the vines are only rain-fed. Certainly, much rain-fed viticulture exists globally, but this is primarily confined to regions that have high seasonal rainfall and/or cool growing season conditions with relatively low evaporative demand, i.e. low leaf-to-air vapor pressure deficits (VPD) and reference evapotranspiration (ET_0_), for example, many European vineyards outside the Mediterranean basin. The prevalence of rain-fed viticulture in semi-arid or arid regions is far less.

Adaptation to drought through exposure of plants to multiple drought events has been reported to have a lasting “learning” effect, leading to a faster response when exposed to the same stress (Bruce et al., 2007). Short term (up to one year) water stress in agricultural crops has been extensively studied, but long-term drought studies have mostly been done in forest tree species (Allen et al., 2010). The underlying reasons for the long term adaptation of plants is still unknown, but theories involving the accumulation of signaling proteins and transcription factors, epigenetics (Tricker et al., 2013), or changes in gene expression have been suggested (Bruce et al., 2007). The long-term adaptation responses of field-grown grapevines to drought have rarely been studied, motivating the current study.

Dry-farming offers an opportunity for viticulturists to sustainably manage their vineyards under conditions of reduced freshwater availability, however, it remains unknown whether dry-farmed vines that have adapted to different levels of soil moisture over long periods perform differently, for example, show varying degrees of water use efficiency. In this context, we conducted a field study on Cabernet Sauvignon grapevines with the hypothesis that dry-farmed vines that had access to lower soil moisture over decadal time scales would be physiologically superior with higher water use efficiency compared to vines that had access to higher soil moisture. This study sheds light on the potential to use dry-farmed grapevines that have potentially adapted to drought conditions as selections for breeding or propagation purposes where an enhanced WUE phenotype is desired.

## 2. Materials and Methods

### 2.1. Vineyard site, climate, plant material

The study was conducted in a commercial vineyard in the Coonawarra region of South Australia (37.28°S, 140.85°E). The Coonawarra region has a long term mean January temperature of 19.3 °C and growing degree days (base 10 °C) of 1511 (Hall and Jones, 2010). The region has a Mediterranean climate with winter rainfall accounting for a majority of annual precipitation (annual rainfall averaging 569 mm (Bureau of Meteorology: http://www.bom.gov.au/climate/averages/tables/cw_026091.shtml); summers are generally warm and dry with droughts not uncommon (Longbottom et al., 2011). Supplemental irrigation is typically required between December and March.

The 8.16 ha vineyard was planted in 1954 to pre-clonal own-rooted *Vitis vinifera* L. cv. Cabernet Sauvignon across 40 rows with a spacing of 3.0 × 1.8 m (row × vine) resulting in a vine density of 1,852 vines ha^−1^ (772 vines ac^−1^). The soil type in this block was “Terrarossa”, a porous and well-drained red clay loam overlaying a limestone sub-layer (Longbottom et al., 2011). The vines were trained to a bilateral cordon at 1.3 m, and spur pruned by hand to approx. seven two-node spurs per linear meter of cordon. No shoot positioning was done during the growing season and, as such, the canopy grew as a sprawl. The entire block was unirrigated (dry-farmed). Vineyard management and integrated pest management practices were applied as per standard practice in the region.

Based on a topography map, 16 vines were selected across the block that visually exhibited relatively high canopy vigour during the previous growing season. The rooting depth, typically the depth to limestone sub-layer in this region, was measured next to each vine approx. 30 cm from the trunk and in the vine row using a two-stroke petrol drill (Model BT45, Stihl, Waiblingen, Germany) fitted with a stainless steel auger (1.5 m length, 22 mm nominal diameter), and tape measure. The 16 vines were then classified into two groups of eight vines based on rooting depth: shallow (SR; rooting depth < 50 cm; average rooting depth = 28.5 cm; range: 14-45 cm) and deep (DR; rooting depth > 50 cm, average rooting depth = 78.1 cm; range = 52-90 cm) rooted.

### 2.2. Canopy light measurements

Leaf area index (LAI) and canopy porosity were measured with a ceptometer (AccuPAR Model LP-80, METER, Pullman, WA, USA) on February 23, 2019 at approximately véraison. One measurement per cordon, two per vine, was taken on the eastern side of the vine. Measurements were made of incident radiation (above the canopy) and light interception in the canopy interior by placing the linear probe horizontally along the cordons on either side of the trunk. Light readings were taken at solar noon (1300 h to 1400 h) when incident light levels were greater than 1500 μE m^−2^ s^−1^.

### 2.3. Soil and vine water status

Pre-dawn leaf water potential (Ψ_pd_) was measured with a Scholander-type pressure chamber (Model 1505, PMS Instruments, Albany, NY, USA) on two leaves per vine, one from each cordon, on March 7, 2019, 22 days after the last rain event, between 0400-0600 h. Ψ_pd_ provides an estimate of soil matric potential or water status (Pagay et al., 2016).

Vine water status was characterized through measurements of stem water potential (Ψ_s_). Two leaves per vine, one per cordon, were bagged in an opaque aluminum foil bag for at least one hour prior to the measurements to stop transpiration and allow leaf equilibration with the shoot (Begg and Turner, 1970). The same pressure chamber was used for these measurements on March 7, 2019, 22 days after the last rain event, between 1200-1400 h.

### 2.4. Leaf gas exchange

Gas exchange measurements were performed on two fully-expanded, healthy leaves per vine, one leaf per cordon, using an open system infrared gas analyzer (IRGA; LI-6400XT, LI-COR Biosciences Inc., Lincoln, NE, USA) on the same day and time as the stem water potential measurements. An external LED light source (LI-6400-02B) attached to the 6 cm^2^ cuvette was used at a fixed PAR value of 1500 μmol m^−2^ s^−1^. The flow rate was set 400 μmol s^−1^ and the reference CO_2_ was set at 400 ppm. Leaves used for gas exchange measurements were selected adjacent to the leaves used for stem water potential measurements.

### 2.5. Yield components

All 16 grapevines were hand harvested on April 1, 2019, approx. one week prior to commercial harvest. Prior to harvest, 200 berries per vine were randomly sampled across the vine for berry weight assessment the same day. The berries were subsequently stored at −18 °C until further analysis of fruit composition. Vine yield and number of clusters were measured on the harvest day, and from this data, average cluster weight was calculated. Average number of berries per cluster was calculated as the ratio of average cluster weight and average berry weight.

### 2.6. Berry carbon isotope analysis

A modified protocol from Gaudillere et al. (2002) was used where 50 berries were randomly selected from the 200 berry sample and hand crushed. The extracted juice was immediately frozen and kept in a −18°C freezer. For the analysis, the juice was defrosted at room temperature, then heated to 70°C and centrifuged to re-solubilise the precipitated acids. Two mL of juice was gently pushed through a one mL column packed with ion exchange resin (Bio-Rad; Mixed bed resin 272 AG501-X8) and then freeze-dried. Approx. 0.8 mg of sample was weighed into a tin capsule and the ratio of carbon isotopes, C^12^ and C^13^ (δ^13^C) was analysed in a NuHorizon Continuous Flow Isotope Ratio Mass Spectrometer (IRMS; Nu Instruments, 274 Wrexham, UK). Eleven samples of each group (deep and shallow soil) were analysed with eight randomly selected replicates (four from deep soil and four from shallow soil vines).

### 2.7. Pruning weight and crop load

During the dormant season, in June 2019, the vines were hand pruned to approx. seven two-bud spurs per meter of cordon, and the pruned canes were weighed using a portable scale. From the pruning weight data, a key vine balance metric, Ravaz Index, was calculated. The Ravaz Index is the ratio of vine yield and pruning weight (Ravaz, 1903).

### 2.8. Statistical analysis

Data was subject to t-tests at *P* ≤ 0.05 using GraphPad Prism statistical software (v.9.0, GraphPad Software, San Diego, USA).

## 3. Results

### 3.1. Environmental conditions

The measurements were conducted over the 2018-19 growing season in Coonawarra, South Australia. The first measurement, on February 23^rd^ 2019, was almost at the end of véraison, a phenological stage which marks the onset of ripening and colour change in red grape cultivars. The environmental conditions during this time were relatively warm and dry with a maximum daily temperature and VPD of nearly 31 °C and 3.30 kPa, respectively (Table 1).

**Table 1.** Environmental conditions on the measurement days and at harvest in 2019. Data obtained from the Bureau of Meteorology automatic weather station at Coonawarra, SA (Station ID: 26091).

| Parameter | February 23 | March 7 | April 1 |
| --- | --- | --- | --- |
| Phenological stage E-L number* | 80% veraison<br>35 | Post-veraison<br>36 | Harvest<br>38 |
| $T_{\max}$ (°C) | 30.7 | 23.5 | 18.6 |
| $VPD_{\max}$ (kPa) | 3.30 | 2.12 | 1.06 |
| $PAR_{\max}$ (W m<sup>-2</sup>) | 601.9 | 601.2 | 66.6 |
| $ET_0$ (mm day<sup>-1</sup>) | 3.25 | 3.39 | 1.38 |

\* Modified Eichorn-Lorenz (E-L) numbering of grapevine phenological stages (Coombe, 1995)

In terms of reference evapotranspiration (ET_0_), approx. 3.3 mm of water was removed from the vineyard over a 24 h period (not including the vines), which was a reflection of the high PAR values (max. 602 W m^−2^). The second measurement day, March 7, was marked by more moderate conditions with daily maximum temperature and VPD of nearly 24 °C and 2.12 kPa, respectively. The ET_0_ that day was approx. 3.4 mm, while the maximum PAR value was 601 W m^−2^, both similar to the first measurement day in February. Harvest occurred on April 1, which was marked by cool and cloudy conditions; the daily maximum temperature and VPD were approx. 19 °C and 1.1 kPa, respectively. The maximum solar radiation that day was a mere 67 W m^−2^ while the ET_0_ was 1.4 mm over a 24 h period.

### 3.2. Canopy light interception and size

Canopy light interception was measured on the sentinel vines using a handheld ceptometer on February 23^rd^, 2019, approx. late-veraison. Leaf area index (LAI) was determined from the light interception measurements using the inversion model developed by Norman and Jarvis (1975); this model relies on measurements or estimates of fraction of transmitted light, solar zenith angle, beam fraction, and extinction angle (amount of light absorbed by the canopy) based on a known leaf angle distribution. LAI values measured in the two classes of vines, shallow rooted (SR) and deep rooted (DR), were statistically indistinguishable; the SR vines had an average LAI value of 5.60 while the DR vines had a LAI value of 5.62 (Fig. 1a).

**Fig. 1.**
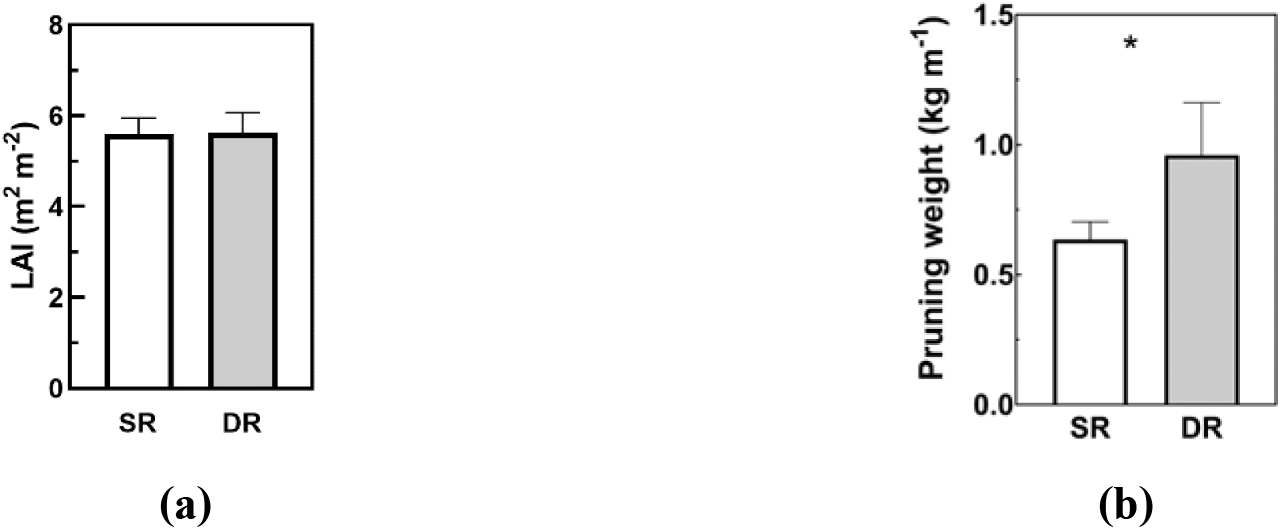
Canopy leaf area index (a) and normalised pruning weight **(b)** of the shallow rooted (SR) and deep rooted (DR) non-irrigated Cabernet sauvignon grapevines on 23/02/2019. Bars represent the standard error of mean (n=8). Significances: *, P ≤ 0.05; **, P ≤ 0.01; ***, P ≤ 0.001, based on Tukey’s HSD test.

During the dormant season, Winter 2019, the vines were hand pruned to two-node spurs and the weight of cane prunings were obtained as an indication of vine size or vigour. The two classes of vines, SR and DR, had significantly different pruning weights (Fig. 1b): SR vines had an average pruning weight of 0.64 ± 0.1 kg while the DR vines had an average pruning weight of 0.96 ± 0.2 kg (*P* = 0.0106).

### 3.3. Soil and vine water status

Predawn leaf water potential measurements (Ψ_pd_) of vines in the two rooting classes, SR and DR, revealed significant differences between the two groups (Fig. 2a). The SR vines had an average Ψ_pd_ of −0.25 ± 0.03 MPa while the DR vines had an average Ψ_pd_ of −0.15 ± 0.01 MPa (*P* = 0.0088). The vine water status as given by the measurement of Ψ_s_ at solar noon indicated that the SR vines had an average value of −0.89 ± 0.04 MPa while the DR vines had an average value of −0.81 ± 0.04 MPa (Fig. 2b). The two groups had similar values of Ψ_s_, i.e. there was no statistically significant difference.

**Fig. 2.**
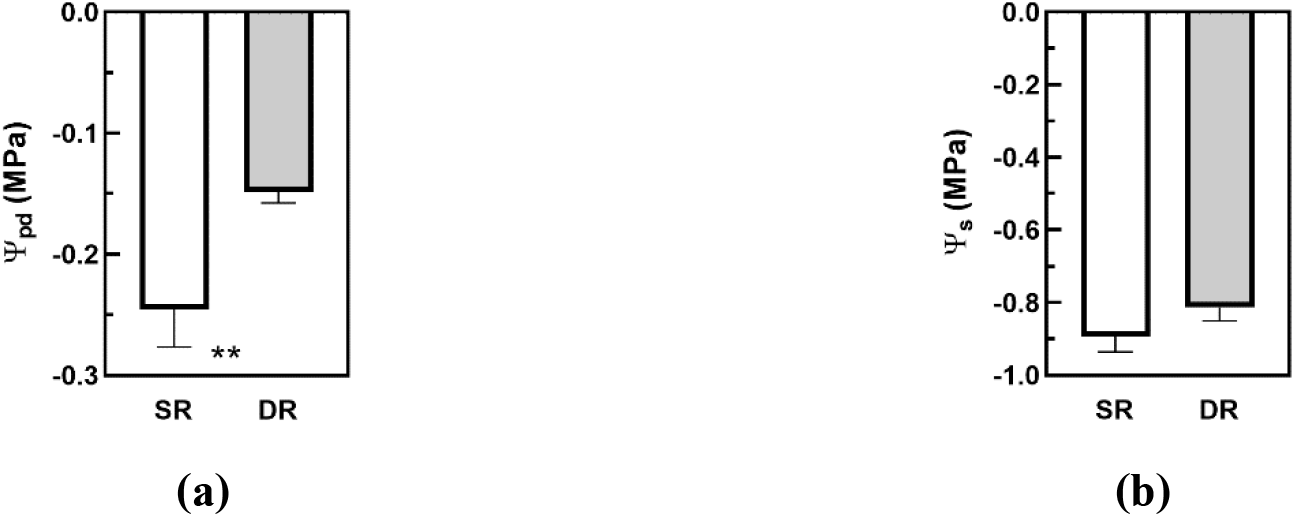
Predawn leaf **(a)** and midday stem water potential **(b)** of the shallow rooted (SR) and deep rooted (DR) non-irrigated Cabernet sauvignon grapevines on 07/03/2019. Bars represent the standard error of mean (n=8). Significances: *, P ≤ 0.05; **, P ≤ 0.01; ***, P ≤ 0.001, based on Tukey’s HSD test.

### 3.4. Leaf gas exchange

Leaf gas exchange was measured using an IRGA and two parameters were obtained for the two rooting classes: leaf stomatal conductance (*g*_*s*_) and net CO_2_ assimilation rate (*A*_*n*_). Intrinsic water use efficiency (*WUE*_*i*_) was calculated as the ratio of *A*_*n*_ and *g*_*s*_. Leaf gas exchange rates were generally observed to be higher in the DR vines compared to the SR vines (Fig. 3a,b). DR vines had an average *g*_*s*_ value of 104 ± 8 mmol H_2_O m^−2^ s^−1^ while SR vines had a significantly lower average value of 74 ± 9 H_2_O m^−2^ s^−1^ (*P* = 0.0333; Fig. 3a).

**Fig. 3.**
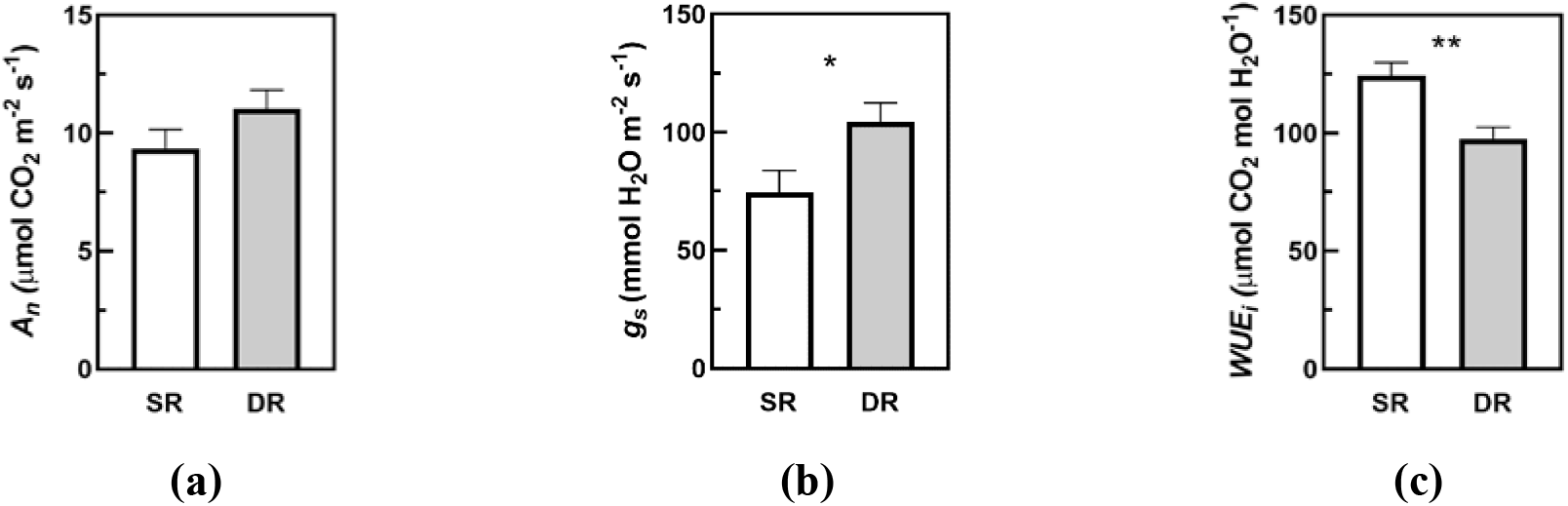
Leaf gas exchange of the shallow rooted (SR) and deep rooted (DR) non-irrigated Cabernet sauvignon grapevines on 07/03/2019. **(a)** stomatal conductance (*g*_*s*_); **(b)** net CO_2_ assimilation (*A*_*n*_); **(c)** intrinsic water use efficiency (*WUE*_*i*_ = *A*_*n*_/*g*_*s*_). Bars represent the standard error of mean (n=8). Significances: *, *P* ≤ 0.05; **, *P* ≤ 0.01; ***, *P* ≤ 0.001, based on Tukey’s HSD test.

Leaf net photosynthesis rates (*A*_*n*_) were observed to be similar between the two classes, SR an DR. The SR vines had average *A*_*n*_ value of 9.4 μmol CO_2_ m^−2^ s^−1^ while the DR vines had an average *A*_*n*_ value of 11.0 μmol CO_2_ m^−2^ s^−1^ (Fig. 3b). As a result of these gas exchange trends, the *WUE*_*i*_ of the two classes were found to be different with the SR vines having higher *WUE*_*i*_ than the DR vines. SR vines had an average *WUE*_*i*_ of 124 ± 6 μmol CO_2_ mol H_2_O^−1^ while the DR vines had a lower average value of 98 ± 5 μmol CO_2_ mol H_2_O^−1^ (*P* = 0.0046; Fig. 3c).

### 3.5. Yield components and crop load

All the sentinel vines were hand harvest on April 1, 2019, and their yield components were measured (Table 2).

**Table 2.** Yield components and crop load (Ravaz Index) of the shallow and deep rooted non-irrigated Cabernet Sauvignon grapevines, 2019. Values are means ± standard error of mean (*n*=8). Significances highlighted in bold indicate *P* ≤ 0.05.

| | Shallow-rooted | Deep-rooted | Significance<br>($P$-value) |
| --- | --- | --- | --- |
| Yield ($\text{kg m}^{-1}$) | $2.48 \pm 0.33$ | $3.75 \pm 0.29$ | **0.014** |
| Average Cluster Number ($\text{m}^{-1}$) | $45.6 \pm 4.4$ | $70.6 \pm 9.1$ | **0.027** |
| Average Cluster Weight (g) | $48.0 \pm 3.7$ | $47.5 \pm 5.3$ | 0.939 |
| Average Berry Weight (g) | $0.90 \pm 0.03$ | $0.89 \pm 0.05$ | 0.904 |
| Average Berries per Cluster | $53.3 \pm 3.8$ | $52.1 \pm 4.3$ | 0.848 |
| Ravaz Index | $3.83 \pm 0.77$ | $4.31 \pm 0.93$ | 0.698 |

SR vines had an average yield of 2.5 ± 0.3 kg m^−1^, which was lower than the average yield of DR vines at 3.8 ± 0.3 kg m^−1^ (*P* = 0.014). Vine yields and cluster numbers were measured on a linear distance of cordon basis due to the high variability of vine (cordon) length in the block resulting from fungal diseases of wood. The average number of clusters per linear distance of cordon was also significantly lower in SR vines compared to DR vines. SR vines had, on average, approx. 46 clusters m^−1^ while DR vines had approx. 71 clusters m^−1^ (*P* = 0.027). The average cluster weights of these two classes of vines were very similar, averaging approx. 48 g. The berry weights were also similar, averaging approx. 0.9 ± 0.03 g in both groups. The number of berries per cluster, an indication of the level of fruit set, was similar between classes, averaging 53 ± 4 g and 52 ± 4 g in the SR and DR vines, respectively. Lastly, crop load as indicated by the Ravaz Index, the ratio of yield and pruning weight, was assessed in the two groups and found to be similar; the SR vines averaged 3.8 ± 0.8 kg kg^−1^ while the DR averaged 4.3 ± 0.9 kg kg^−1^.

### 3.6. Fruit carbon isotope ratio

The carbon isotope ratio (*δ*^13^*C*) of the berry samples harvested from the two rooting classes had very similar values: SR had an average *δ*^13^*C* value of −25.0 ± 0.2 ‰ while the DR fruit had an average *δ*^13^*C* value of −25.4 ± 0.2 ‰ (Fig. 4). The two rooting classes, DR and SR, were statistically indistinguishable in terms of their fruit *δ*^13^*C* values, which indicates that their seasonal integrated water use efficiency was similar.

**Fig. 4.**
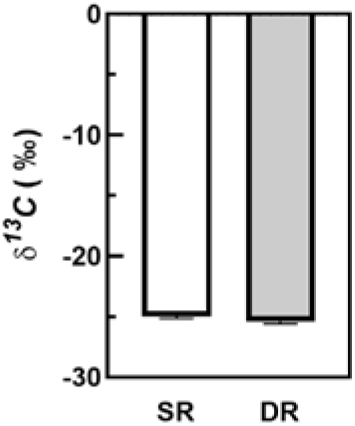
Carbon isotope (*δ*^13^*C*) values of fruit harvested from shallow rooted (SR) and deep rooted (DR) non-irrigated Cabernet sauvignon grapevines. Bars represent the standard error of mean (*n*=8).

## 4. Discussion

Grapevines are able to respond and adapt to both short and long term changes in environmental conditions such as soil water deficits through mechanisms such as stomatal closure (Chaves et al., 2010; Pagay et al., 2016), osmotic adjustment with compatible solutes such as proline and sorbitol (During, 1984; Rodrigues et al., 1993) as well as potassium and calcium (Degu et al., 2019), non-photochemical quenching (of chlorophyll), activation of the reactive oxygen species (ROS) detoxification pathway (antioxidant system) (Beis and Patakas, 2012), and hydraulic vulnerability segmentation (Schultz, 2003; Shelden et al., 2017). Variations in soil moisture across individual vineyard blocks have been documented, primarily in the context of *terroir* studies. These variations have been attributed to soil type (Brillante et al., 2016; Willwerth and Reynolds, 2020) and soil depth (Smart et al., 2006; Wheaton et al., 2008), both which have been linked to grapevine growth and productivity, i.e. yield, and grape and wine chemical composition (Renouf et al., 2010; Storchi et al., 2005).

### 4.1. How do soil structure and depth influence vine performance?

Terrarossa soils, which dominate Coonawarra vineyards, are highly porous clay loams with iron oxide overlaying limestone at variable depths from 50 to 150 cm (Longbottom et al., 2011). The variations in soil depth, constraining vine rooting depth, and the porous nature of Terrarossa soils are significant factors influencing soil moisture availability and consequently vine water and nutrient status. Our study observed variations in soil depth across an 8 ha vineyard block ranging from 14 cm to 90 cm. It is known that soil texture plays a role in determining vine rooting depth with coarse soils allowing for greater rooting depths while, by contrast, more restrictive heavier clay-based soils that restrict root growth contributing to greater vine water stress and lower yield (van Leeuwen et al., 2004). Furthermore, clay-based soils can achieve a degree of porosity with adequate humus and calcium (Seguin, 1986), the latter being a key component of the limestone sub-strata of Coonawarra soils (Longbottom et al., 2011). Vines in the present study were not planted in different soil types, but rather in varying soil depth; vine responses in shallow soils in the present study were similar to those reported by van Leeuwen and colleagues in predominantly clay soils (van Leeuwen et al., 2004). In our study, vines in shallow soils (SR) had lower vine size (pruning weight), soil moisture (Ψ_pd_), leaf gas exchange, and yield compared to those in deeper soils (DR). Notably, the lower soil moisture in the shallow-rooted vines did not translate to lower vine water status (Ψ_s_) or net photosynthesis (*A*_*n*_), which, in conjunction with lower *g*_*s*_, resulted in higher *WUE*_*i*_ (Fig. 3). The reduction of *g*_*s*_ due to lower soil moisture could be attributable to root-to-shoot chemical signals, e.g. abscisic acid (Correia et al., 1995; Gowing et al., 1990; Stoll et al., 2000) and, more recently, small peptides (Takahashi et al., 2018). These peptides, particularly those belonging to the CLAVATA3 family, have been shown in *Arabidopsis* to be a root-to-shoot signal inducing abscisic acid (ABA) biosynthesis in leaves inducing stomatal closure and enhanced drought tolerance (Takahashi et al., 2018). In addition to chemical signaling, lower *g*_*s*_ may also be the result of hydraulic signaling, e.g. changes in plant hydraulic conductivity (Dayer et al., 2020).

The decrease in *g*_*s*_ while maintaining *A*_*n*_ (hence increasing *WUE*_*i*_) could be an adaptation response of grapevines to the typically high VPD of xeric environments (Maroco et al., 1997). The lack of change of *A*_*n*_ in response to varying soil moisture could be due to the relatively mild degree of vine water stress observed in this study. These similar *A*_*n*_ values could also be due to its lower sensitivity than *g*_*s*_ to moderate soil moisture deficits (Chaves et al., 2010). In the present study, maintenance of leaf photosynthesis, which is quite resistant to water stress (Flexas and Medrano, 2002), despite lower leaf conductance in shallow-rooted vines suggests that non-stomatal (or biochemical) factors, including maximum rates of electron transport (*J*_*max*_) and carboxylation (Vc_max_), and mesophyll conductance (*g*_*m*_) may be similar between the two groups of vines. This hypothesis could form the basis of a follow on study.

The lack of differences in *A*_*n*_ between soil depths could explain the similar berry weights observed (Table 2). However, it was surprising to observe a lack of difference in water use efficiency as quantified by *δ*^13^*C*. This parameter is a measure of seasonal vine water use efficiency between fruit set and harvest as it integrates the discrimination ability of the leaf photosynthetic apparatus to carbon isotopes, ^12^C and ^13^C, in the atmosphere and can be used to quantify differences in soil moisture availability (Gaudillere et al., 2002). The lack of differences in *δ*^13^*C* could relate to the narrow range of soil and/or vine water status values observed in this study, particularly between véraison and harvest.

### 4.2. Is there evidence of drought stress priming?

Seasonal droughts are a common feature of Mediterranean climate, which are characterized by warm, dry summers and wet winters. Our test site provided a unique situation to evaluate the long-term (> 65 years) drought adaptation potential of field-grown grapevines under different levels of soil moisture availability in a Mediterranean climate with seasonal drought. The ability of shallow-rooted grapevines to maintain superior vine water status, leaf gas exchange and berry size despite having lower soil moisture availability in the present study indicates a degree of long-term drought adaptation of those vines, a trait that is potential inheritable (Tricker et al., 2013). A study comparing the drought tolerance of progeny of several sub-species of sagebrush (*Artemesia tridentata*) obtained from naturally varying aridity habitats found that those from relatively arid habitats had the highest drought tolerance (Maier et al., 2001). The exposure of plants to multiple drought events, which are not uncommon in South Australian vineyards, have been shown to “prime” the plants and enhancing their sensitivity to future droughts (Bruce et al., 2007).

The impacts of a warming climate are observed across the decades of climate analysis of this trial. Heat units have risen from the initial planting of the vineyard in 1954 to harvest in 2019 by 37% concomitant with a reduction in annual rainfall of 14% (Table S1). An artifact of the trial has identified considerable year to year variability in the last decade than earlier decades, which is consistent with trends of climate change (http://www.bom.gov.au/state-of-the-climate/documents/State-of-the-Climate-2020.pdf).

Evidence of long-term studies on adaptation to multiple drought events in horticultural crops is limited. In forests, drought events lead to tree mortality through water stress, carbon starvation, and biotic stresses (Allen et al., 2010). Long-term drought responses, e.g. structural and biomass changes, and tree mortality were not predictable from short-term drought responses, e.g. leaf gas exchange and hydraulics, in a tropical rainforest over 15 years (Maier et al., 2001). Our short-term observations of field-grown grapevines provided a unique insight into their adaptation potential to cycles of drought over decadal timescales. The adaptation of shallow-rooted grapevines to cycles of drought over long timescales could relate to epigenetic modifications of DNA as observed in other crops including wheat (Budak et al., 2015), rice (Wang et al., 2011), and tomato (Benoit et al., 2019). The possible ‘stress priming’ of grapevine individuals in shallow soils could enhance their resilience to future drought events (Crisp et al., 2016).

The practical significance of this finding is the ability to identify and select potentially superior, drought-tolerant clones within a sub-species that can be propagated for future vineyard plantings. Future work in this area could investigate the roles of non-stomatal responses to photosynthesis and reactive oxygen (ROS) detoxification species, as well as epigenetic changes that may explain the superior phenotype of the shallow-rooted vines.

## 5. Conclusions

Our study unveiled the potential of selecting drought-tolerant grapevines by exploiting their long-term adaptation to soil moisture differences and provides yet another alternative to traditional breeding. Although drought rarely occurs in isolation as an environmental stress in the field, typically co-occurring with heat and light stress and, in many Australian vineyards, salinity stress, dry-grown grapevines appear to have the potential to adapt to these stresses over decadal timescales. The increased ‘fitness’ resulting from this adaptation may confer superior tolerance to other abiotic and biotic stresses, including pests and diseases. In viticulture, climate change, and droughts in particular, will continue to constrain our natural resources, e.g. freshwater, and challenge current viticultural practices, e.g. irrigation management, therefore requiring a more extensive toolkit of genetic resources to better prepare for future drought scenarios.

## Acknowledgements

The authors would like to thank Wynns Coonawarra Estate for access to their vineyard, and for providing material and in-kind support. Tarita Furlan would like to thank Casella Family Brands for financial support through the Casella Family Brands Scholarship.

